# Binocular visual input is essential for prey capture but not attention to prey in the praying mantis *Sphodromantis lineola*

**DOI:** 10.64898/2026.07.14.738220

**Authors:** Emiliano Kalesnik, Théo Robert, Julieta Sztarker, Vivek Nityananda

## Abstract

Binocular vision provides animals with several evolutionary advantages. In praying mantises, one of these advantages is stereopsis that has different effects while attending to prey and capturing prey. However, the importance of binocular, compared to monocular, visual input has not been tested in either stage of predation. We therefore used an insect 3D cinema to present mantises with binocular and monocular stimuli to either prime their attention or elicit prey capture. We used a previous paradigm where a wide-field figure motion cue attracts mantis attention, which leads to predatory responses to a small-field elementary motion target. We found that binocular visual input enhances attention to cues, but monocular cues are also effective. However, prey capture responses were fundamentally dependent on binocular input. Thus, binocularity appears to be fundamental to prey capture in mantises but not for attending to prey.

## Introduction

Binocular vision is widespread across the animal kingdom and can provide an animal with several evolutionary advantages [1]. The most well studied of these is stereopsis or stereo vision [2]. Stereopsis involves visual information from both eyes being compared to calculate the distance to an object. The difference in object position in each eye is referred to as visual disparity. Greater disparities indicate nearer objects, and lower disparities indicate objects further away. However, binocular vision could provide other advantages [1]. Having two eyes, could for example, allow for redundancy, in case one eye was injured. It could also increase sensitivity in the region of binocular overlap and enable seeing around obstacles.

Several insects use binocular vision for adaptive behaviour in different contexts. Bees compare optic flow between both eyes to stabilize flight [3]. The separation between the eyes of stalk-eyed flies could help them see around obstacles in cluttered environments [4]. Insect predators could have special advantages, given that they often have frontal facing eyes with high binocular overlap. Damselflies, for example, combine motion information from both eyes for target pursuit [5]. Praying mantises also use binocular vision to track and capture prey [6,7]. Importantly, they are the only insects known to have stereopsis [8,9].

Stereopsis in mantises has recently been studied using an insect 3D cinema [9,10]. Mantises were fitted with colour filters on their eyes that enabled the presentation of different virtual stimuli to each eye (Fig. 1A). The set-up was used to vary the disparity of stimuli and simulate objects at different depths. Mantises in this set-up strike at prey where the two visual lines of sight cross (crossed disparity, Fig 2A) to create interocular disparity simulating prey that are nearby (2.5 cm) but not at prey simulated to be farther away (10 cm, zero disparity, Fig. 3A). They also do not strike at prey when the same visual stimuli are swapped in each eye, so that the lines of sight do not cross, leading to an impossible target (uncrossed disparity). This approach has revealed that, unlike in primates, stereopsis in mantises relies on comparing the disparity between regions that have motion or other temporal changes in each eye [11].

**Figure 1:**
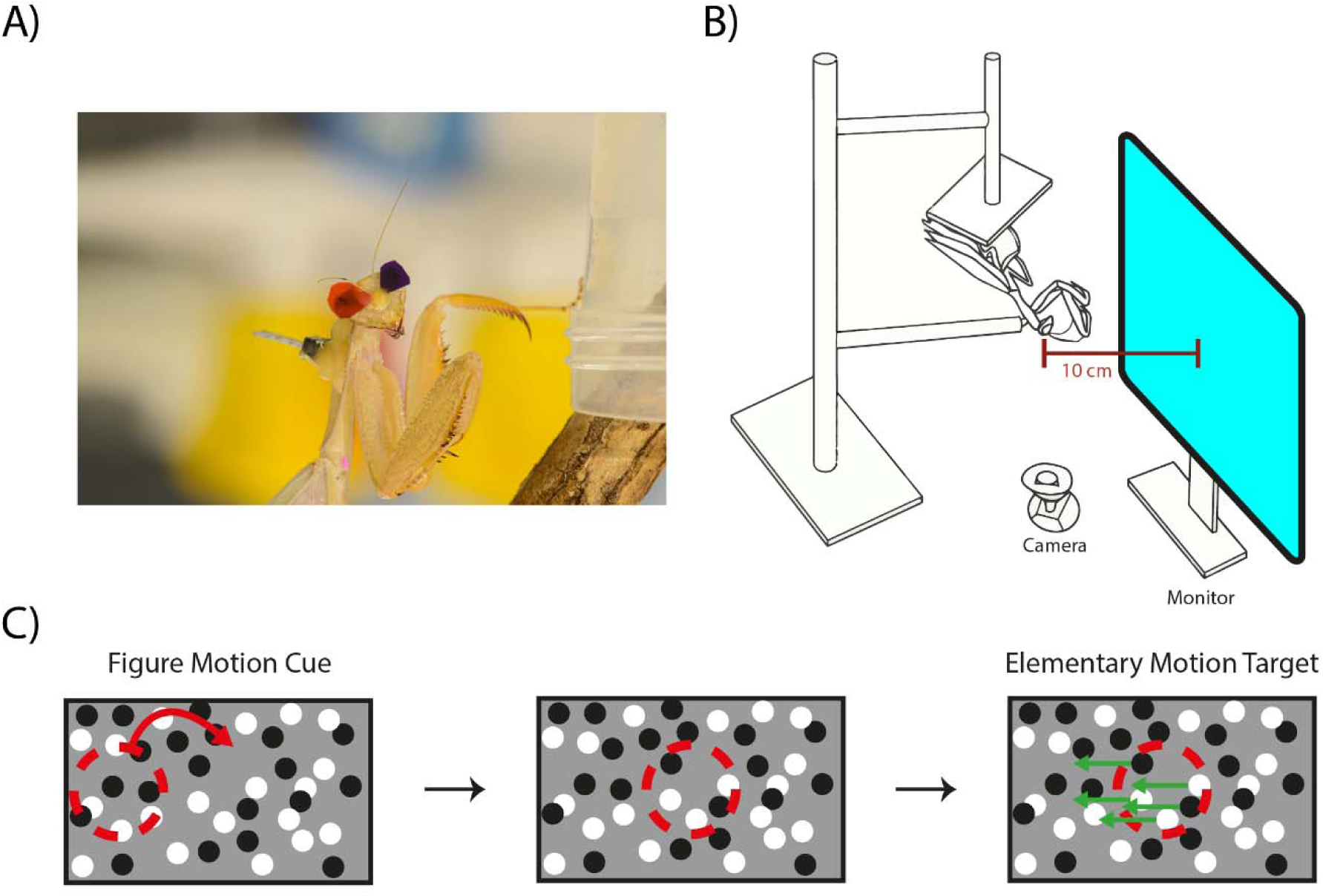
Experimental set up and stimuli. A) Mantis with 3D glasses and two-pin header on the back. Photo credit: Hugo Le Lay, B) Experimental set-up, C) Schematic depiction of the stimuli: the left panel depicts the drift-balanced spiral figure motion cue (red dotted circle); the centre panel represents the inter-stimulus interval (2 s) where there is no motion; and the right panel represents the small-field elementary motion target (small dots moving within a restricted area shown by the red dotted circle).

**Figure 2:**
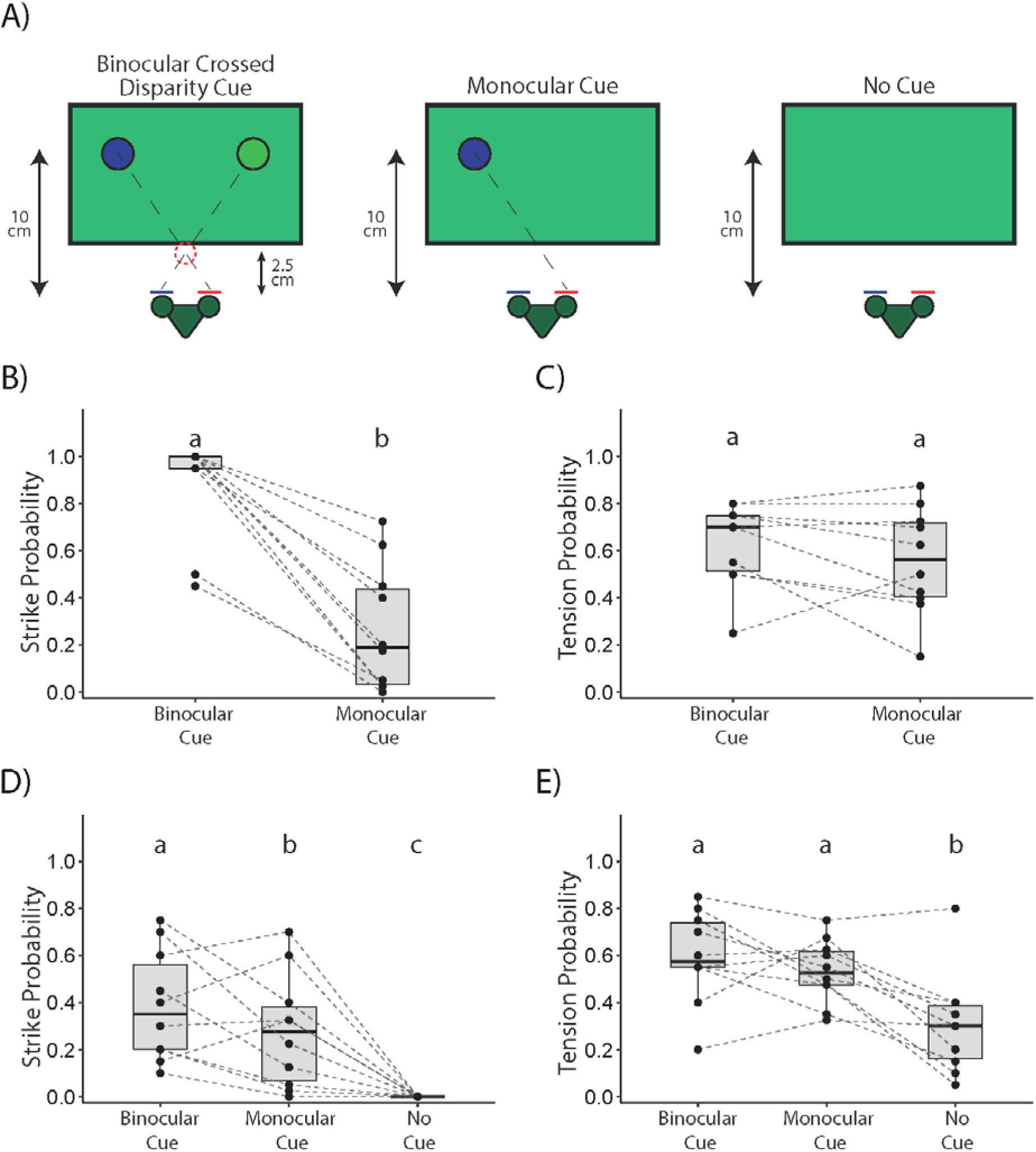
Responses in Experiment 1. A) Schematic depiction of the different conditions of the Figure Motion cue. Left panel depicts binocular crossed disparity with a simulated target 2.5 cm from the mantis. Centre panel depicts a stimulus presented to a single eye. Only one colour is depicted here but across trials both colours were presented equally as monocular stimuli. Right panel depicts the condition with no cue. B) Probability of strikes to the Figure Motion Cue, C) Probability of tensions to the Figure Motion Cue, D) Probability of strikes to the Elementary Motion target following different conditions of the Figure Motion cue, E) Probability of tensions to the Elementary Motion target following different conditions of the Figure Motion cue. Different letters indicate significant differences (P < 0.05).

**Figure 3:**
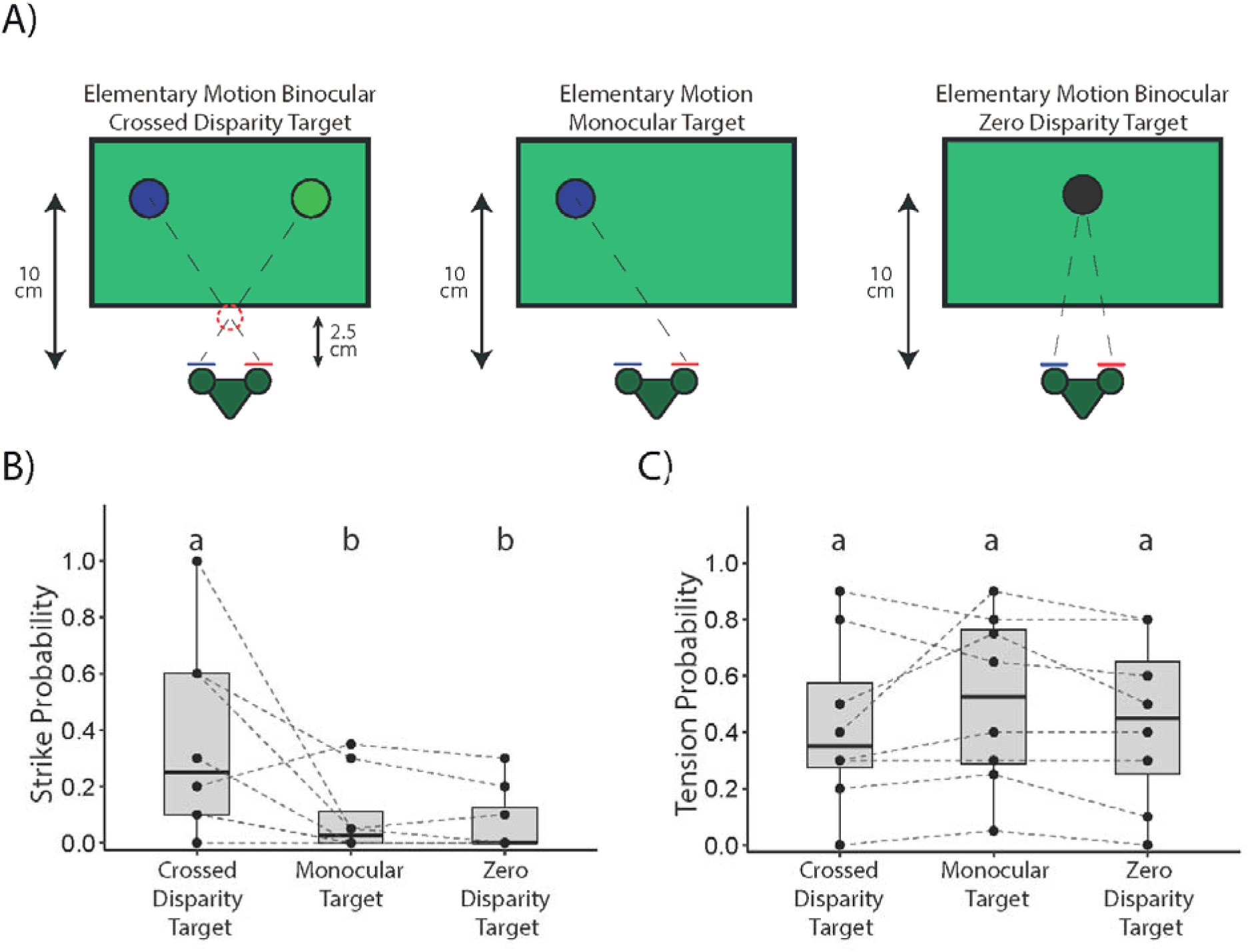
Responses in Experiment 2. A) Schematic depiction of the Elementary Motion conditions. Left panel depicts binocular crossed disparity with a simulated target 2.5 cm from the mantis. Centre panel depicts a target stimulus presented to a single eye. Only one colour is depicted here but across trials both colours were presented equally as monocular stimuli. Right panel depicts a binocular target with zero disparity. Note that the actual stimulus consisted of smaller dots with movement in a limited region (Fig. 1C left panel) B) Probability of strikes during the Elementary Motion, C) Probability of tensions during the Elementary Motion. Different letters indicate significant differences (P < 0.05)

Prey capture in mantises is thus facilitated by motion-dependent stereopsis, a key advantage provided by binocular vision. Several types of motion can be used by mantises for stereopsis. This includes both second-order figure motion and small-field elementary motion [12]. Second-order figure motion involves wide-field motion only detected by correlating higher order motion statistics (Fig. 1C, Left Panel), while small-field elementary motion consists of localized movements detected by elementary motion detectors (Fig. 1C, Right panel). Responses to elementary motion are dependent on stereo disparity, with mantises striking at crossed but not uncrossed disparities. However, mantises will not strike at this form of motion unless it is first preceded by wide-field figure motion (Fig. 1C), suggesting a form of attentional cuing. The attentional cuing effect of figure motion, however, occurs for both crossed and uncrossed disparities of figure motion.

This suggests that there are two different but interconnected systems underlying predation which we can characterize as attending to prey and prey capture, with the latter being more strongly dependent on stereo disparity. It is still unclear, however, how important binocular vision is to both these systems. The cuing effect of figure motion is stronger for crossed stereo disparities simulating a target at 2.5 cm but it is still effective for other disparities, including uncrossed disparities [12]. All these conditions however, provided binocular visual input and providing only monocular input could remove this attentional effect. Responses to elementary motion have also only been tested with binocular stimuli and so far, mantises have not been tested with monocular virtual stimuli, or small-field stimuli simulated to be at real depths that are too far to capture. Comparing responses to binocular and monocular virtual stimuli would thus reveal which aspects of predation rely primarily on binocular input and where monocular input can also be effective.

To test mantis responses to these different stimuli, we presented mantises with virtual stimuli consisting of a wide-field figure motion stimulus (the cue; Fig. 1C, left panel) followed by small-field elementary motion of dots (the target; Fig. 1C, right panel). Taking advantage of the insect 3D cinema where different stimuli can be presented to each eye, we presented stimuli to either one eye or both, thus providing either binocular or monocular stimulation. In separate experiments, we tested the effectiveness of binocular and monocular stimuli in the figure and elementary motion phases. We hypothesised that binocular stimuli would be more effective as attentional cues than monocular stimuli, eliciting stronger responses to the subsequent prey. Given that prey capture is especially dependent on stereo disparity, we also hypothesised that mantises would be more likely to make predatory responses to binocular stimuli simulated to be nearby compared to monocular stimuli or stimuli simulated to be farther away.

## Methods

### Experimental subjects

All experiments were conducted on adult female mantises of the species *Sphodromantis lineola* maintained in a culture at Newcastle University. Mantises were housed individually in semi-transparent cages (13 x 13 x 20 cm) in a climate-controlled environment maintained at a temperature of 25 °C. Mantises were fed only 1 cricket per week on the weekends throughout the duration of the experiments to maximize predatory behaviour. Experiments were conducted in a within-subject design, so all mantises experienced all experimental conditions.

### Preparing and fixing the 3D glasses

Red and purple colour filters (LEE filters 135 Deep Golden Amber and 797 Purple respectively) were used to deliver anaglyph 3D images [10] (Fig. 1A). These filters achieved separate presentation of images to each eye based on their different spectral content, enabling the delivery of monocular or binocular content as well as manipulation of stereoscopic disparity. The transmittance spectra of these filters were as previously published [10]. Tear-drop-shaped glasses were cut out of the filters with a maximum diameter of 1 cm. Mantises were pinned down using modelling clay, and the glasses were affixed to the front of the mantises using beeswax applied with a wax melter (Denta Star S ST 08) so that a different colour filter covered each eye (Fig. 1A). The assignment of colour filters (red and purple) to the left and right eyes was counterbalanced across mantises.

### Visual stimulation and experiments

Mantises were fixed to a stand with an electrical pin header of two pins fitted onto their backs with beeswax (Fig. 1A) that attached to its counterpart on the testing stand. This kept the mantises at a fixed distance of 10 cm from a computer monitor. Mantises were fixed upside down on the stand and allowed to hold onto a moveable cardboard disk with their feet, but were able to move their head freely, as well as make strikes with their forearms. The mantises were filmed during the experiments with a Kinobo USB B3 HD Webcam (Point Set Digital Ltd) placed below the screen (Fig. 1B).

Visual stimulation was provided at a frame rate of 60 Hz on a DELL U2413 LED monitor (1,920 × 1,200 pixels; 51.8 × 32.4 cm). The background for all stimuli consisted of blue and green dots against a cyan background. The dots had a diameter of 25 pixels (1.8°) and a density of 55 dots in every 100 × 100 pixel square: 50% of the dots at the maximum luminance (“white” dots) and 50% at the minimum luminance (“black” dots). Through the glasses, each eye would be able to see only blue or green dots as dark and light dots against a “gray” background (schematically depicted in Fig 1C). The dots were uncorrelated across the 2 eyes, i.e., each eye saw a completely different dot pattern.

Prior to all experiments, mantises were tested for motivation by using a dark disk that spiralled in from the periphery to the centre. The size and disparity of this disk was chosen to simulate a target 1 cm in diameter and 2.5 cm from the mantis. This is a stimulus that mantises respond to strongly and strike at [9]. Experiments were only carried out if mantises struck at two consecutive presentations of this stimulus. Similarly, the data from an experiment were excluded if the mantis did not meet the same criterion after the experiment was carried out.

Subsequently in all experiments, mantises were presented with a wide-field figure motion “cue” followed by a small-field elementary motion “target” stimulus, separated by a 2 s pause (Fig. 1C). The figure motion consisted of a “drift-balanced” disc stimulus that spiralled into the centre of the screen from the periphery in the same movement as the motivational stimulus. The disc here consisted of notional disc with dots that differed from the background, but without any motion. This means that it was effectively a moving hole revealing a different pattern of dots behind those displayed on the screen. This stimulus thus had no elementary motion. The subsequent localized, elementary motion target stimulus consisted of a stationary region in the centre of the screen within which dots moved from one edge of the region to the other with the dots that disappeared being replaced by newly generated ones on the initial edge. This stimulus thus had no second-order figure motion but had elementary motion.

The question and analysis for Experiment 1 was preregistered at aspredicted.com (https://aspredicted.org/n4z5s6.pdf). In Experiment 1 (n = 10 mantises), the elementary motion target stimulus was fixed at a disparity simulating a depth of 2.5 cm while the figure motion cue varied across different conditions: Binocular (simulating a depth of 2.5 cm), Monocular (seen through the red lens), Monocular (seen through the purple lens) or No Cue. In Experiment 2 (n = 8 mantises), the figure motion cue remained constant as a binocular stimulus with stereoscopic disparity simulating a depth of 2.5 cm. The elementary motion target stimulus was presented in four conditions: it varied between Binocular with crossed disparity (simulating a depth of 2.5 cm), Monocular (seen through the red lens), Monocular (seen through the purple lens) or Binocular with zero disparity (with the stimulus appearing at the distance of the monitor, 10 cm). For both experiments, conditions were presented in interleaved trials presented in randomized order across two experimental runs of 40 trials each. This resulted in 10 replicates per run and a total of 20 replicates per condition for each mantis. The direction of elementary motion (left or right) was counterbalanced across all trials within an experimental run.

### Quantification and Statistical Analysis

All trial recordings were made so that the computer screen was not visible and the experimental condition was masked. These movies were then coded without knowledge of the experimental condition. Two types of behaviours were coded: strikes and tensions. Strikes were rapid extensions of the forelegs to capture prey targets perceived to be nearby, while tensions were preparatory movements for a strike, which was eventually unreleased. In both experiments, an indicator dot was visible on screen in the videos during the figure motion phase and disappeared in the elementary motion phase. This dot was hidden from view of the mantis with a cardboard frame and was used to classify responses separately to figure motion and elementary motion stimuli during video coding.

Since the data from the responses were binomial (response or no response), we used a binomial family and a logit link function in GLMMs (from the R package lme4 [13]) to analyse the data. In all experiments, the response variable was the probability of a predatory response (“yes” or “no”) during the figure and the elementary motion stimuli display. The independent factor was the visual condition of the Figure Motion (Experiment 1) or the Elementary Motion (Experiment 2), and the identity of the animal was included as a random factor. All monocular trials were pooled into a single ‘monocular’ condition, and data were pooled across motion directions after confirming that both of these factors had no significant effect on the response. We looked for the main effect of visual condition in each experiment with three levels. In Experiment 1, the three levels were binocular crossed disparity cue, monocular cue and no cue. In Experiment 2, the three levels were binocular crossed disparity target, monocular target and binocular zero disparity target. All data analyses were performed using R Version 4.3.1 (R Core Team, 2023).

## Results

### Experiment 1: Binocular attentional cues elicit more strikes to subsequent prey

We varied the binocularity of a wide-field figure motion cue and tested the predatory responses of mantises to both the cue and subsequent elementary motion target (Fig. 2A). We measured two types of responses, *strikes* which are rapid extensions of the forelegs to capture prey, and *tensions* which are preparatory movements for a strike, eventually unreleased. Mantises responded to both the cue and the subsequent target.

#### Response to the cue

Compared to monocular cues, mantises were significantly more likely to strike at binocular cues, with crossed disparity simulating a target 2.5 cm from the mantis (GLMM: Estimate = 5.23, P < 2 X 10^−16^; Fig. 2B). Both binocular and monocular cues, however, were equally likely to lead to tensions (GLMM: Estimate = 0.34, P = 0.071; Fig. 2C).

#### Responses to the target

Strike probability was significantly higher following binocular compared to monocular cues (GLMM: Estimate = 0.57, P = 0.015; Fig. 1D). Furthermore, both cue types significantly increased the strike probability relative to the No Cue condition (GLMM, Binocular cue: Estimate = 4.95, P < 0.0001; Monocular cue: Estimate = 4.39, P < 0.0001), where strikes were entirely absent.

A different pattern was observed for tensions. Both binocular and monocular cues were equally effective at leading to tensions to subsequent targets (GLMM: Estimate = 0.27, P = 0.29; Fig. 1E). Notably, there was a low probability of tensions to the target even in the No Cue condition, although this was significantly lower than that observed following either of the two other cues (GLMM, Binocular cue: Estimate = 1.29, P < 0.0001; Monocular cue: Estimate = 1.02, P < 0.0001).

### Experiment 2: Strikes to prey require binocular visual input but tensions do not

We tested for strikes and tensions in response to small-field elementary motion stimuli varying in binocularity and disparity after cuing mantises with crossed disparity figure motion simulating a cue 2.5 cm from the mantis (Fig. 3A). Binocular elementary motion stimuli with crossed disparity simulating a target at 2.5 cm from the mantis were most effective at eliciting strikes (Fig. 3B). Both monocular stimuli and binocular stimuli with zero disparity, which simulated prey 10 cm from the mantis, elicited almost no strikes. Mantises were thus significantly more likely to strike at binocular crossed disparity stimuli compared to monocular or zero disparity stimuli (GLMM: Estimates = 2.10 and 2.37 respectively, P < 0.0001; Fig. 3A). There was no difference in the likelihood of strikes to monocular or zero disparity stimuli (GLMM: Estimate = 0.27, P = 0.87). All three stimuli were, however, equally likely to elicit tensions (GLMM, Crossed disparity - Monocular: Estimate = −0.49, P = 0.29; Crossed Disparity – Zero Disparity: Estimate = - 0.07, P = 0.98; Monocular – Zero Disparity: Estimate = 0.42, P = 0.40; Fig. 3C).

## Discussion

Our results show different effects of binocularity on attentional cuing and prey capture in mantises, as well as on different types of predatory responses. While binocularity enhanced the cuing effect of a wide-field figure motion stimulus on eliciting subsequent strikes, monocular cues were also effective. However, small-field elementary motion stimuli needed to be binocular to lead to predatory strikes. Tensions were increased by attentional cuing by both binocular and monocular figure motion stimuli and were equally likely for binocular elementary motion stimuli of either disparity, and monocular stimuli.

Binocular vision has previously been implicated in mantis distance estimation in the context of predator avoidance [7] or object tracking [6]. In those experiments, monocular stimuli were presented by painting over one eye of the mantis. The changed response could therefore have been due to the different physical conditions experienced by these mantises compared to controls. The use of virtual stimuli and a 3D insect cinema in our experiments avoids this problem, as binocular and monocular stimuli could be presented to, and compared within the same individual. Previous results implicated binocular vision as being important for changes in distance but not for lateral movements. Our results show that even without motion in depth, binocular vision is important for mantises and affects both attention to prey and prey capture.

Binocular input affected strikes and tensions differently. Both binocular attention cuing and local motion elicited more strikes but were not different in terms of eliciting tensions. This indicates that both binocular and monocular stimuli were detected and fed into pathways for predatory responses. The difference between strikes and tensions suggests that binocularity increases motivation to strike, potentially stimulating the prey capture system above a neural threshold. In the case of localized prey motion, binocular visual input appears to be critical for the motivation to strike. Interestingly, attention to prey can be triggered by monocular stimuli even if not to the same level as binocular stimuli. This would support the redundancy hypothesis for binocularity – it would be advantageous for an individual to attend to prey even if visible with only one eye. It is an open question whether and how mantises with one damaged eye might capture prey given the low level of strikes to monocular prey. Potentially, this response might change over time as mantises grew accustomed to purely monocular stimuli.

The importance of stereo disparity in attentional cuing is understudied. Some studies from humans show that disparities simulating nearer objects are more strongly attended to [15,16]. Similar research has rarely been carried out in insects. There is suggestive neural evidence for disparity-dependent top-down visual attention [17]. However, a recent study using a cued selective attention paradigm in mantises found weak evidence for disparity influencing attention [18]. In the paradigm used in this study, there is some previous evidence that disparities simulating nearby targets have a stronger effect on attention, but cues of all tested disparities nonetheless could still had a cuing effect [12]. Our results extend these findings showing that even monocular cues can be effective at cuing attention. The presence of wide-field motion is thus sufficient to cue attention, even without binocular input. Binocular input and stereopsis are therefore not necessary for attracting mantis attention, but might have other effects, including affecting the sensitivity to stimuli and the duration of attention. These would be important areas for further research.

## Acknowledgements

VN and TR were supported by a BBSRC David Phillips Fellowship BB/S009760/1 to VN. EK was supported by the European Union’s Horizon 2020 Research and Innovation Program under the Marie Sklodowska-Curie grant agreement No 778062, ULTRACEPT to JS.

